# Time-dependent Role of Bisphosphonates on Atherosclerotic Plaque Calcification

**DOI:** 10.1101/2022.02.14.479950

**Authors:** Amirala Bakhshian Nik, Hooi Hooi Ng, Manuel Garcia Russo, Francesco Iacoviello, Paul R. Shearing, Sergio Bertazzo, Joshua D. Hutcheson

## Abstract

Atherosclerotic plaque calcification directly contributes to the leading cause of morbidity and mortality by affecting the plaque vulnerability and rupture risk. Small microcalcifications can increase plaque stress and promote rupture, whereas large calcifications can stabilize plaques. Drugs that target bone mineralization may lead to unintended consequences on ectopic plaque calcification and cardiovascular outcomes. Bisphosphonates, common anti-osteoporotic agents, elicited unexpected cardiovascular events in clinical trials. Here, we investigated the role of bisphosphonates treatment and timing on the disruption or promotion of vascular calcification and bone mineral in a mouse model of atherosclerosis. We started the bisphosphonate treatment either before plaque formation, at early plaque formation times associated with the onset of calcification, or at late stages of plaque development. Our data indicate that long term bisphosphonate treatment (beginning prior to plaque development) leads to higher levels of plaque calcification, with a narrower mineral size distribution. When given later in plaque development, we measured a wider distribution of mineral size. These morphological alterations may associate with higher risk of plaque rupture by creating stress foci. Yet, bone mineral density positively correlated with the duration of bisphosphonate treatment.

## 1. Introduction

Atherosclerosis represents the most common cause of cardiovascular disease, and atherosclerotic plaque rupture results in arterial thrombosis, leading to heart attacks and strokes [1, 2]. Vascular smooth muscle cell migration and osteogenic differentiation promote calcification within the atherosclerotic plaque [3]. Plaque calcification occurs in 53% and 32% of American male and female atherosclerotic patients, respectively [4]. Comprehensive analyses of coronary artery calcium score have demonstrated a strong positive correlation between calcification and all-cause mortality, including cardiovascular and coronary artery disease [5]. Though overall calcium score positively predicts cardiovascular morbidity, local effects of calcification on plaque stability is determined by calcification size and morphology.

Recent clinical studies reported percent calcified plaque volume as a key factor for plaque stability [6]. Plaques with high percentages of calcification were more stable and less likely to rupture. Plaque vulnerability classically associates with low collagen content in the fibrous cap, which compromises its tensile strength [7]. *In silico* studies highlight the presence of destabilizing microcalcifications (5-15 μm)—undetectable due to resolution limits of the traditional clinical imaging modalities—in the cap of the vulnerable plaques as a determinant of their biomechanical failure [8, 9]. Large macrocalcification (> 50 μm) stabilizes the plaque through tissue stress factor reduction. However, microcalcifications destabilize the plaque by creating stress foci within the fibrous cap due to large mismatch in material properties between the stiff minerals and the surrounding hyperelastic extracellular matrix [8-12]. Prospective clinical data from the Multi-Ethnic Study of Atherosclerosis (MESA) cohort corroborates the biomechanical model predictions linking microcalcification with plaque rupture [12]. These data suggest that coronary artery calcification volume directly contributes with higher risk of coronary heart disease and cardiovascular events. Interestingly, calcium density inversely correlates with cardiovascular event risk [12].

Patients with bone disorders are also at high-risk for cardiovascular events [13, 14]. Unbalanced bone turnover (either high or low bone formation rate) alters the serum calcium and phosphate levels and increases the risk of vascular calcification [15]. Bisphosphonates (BiPs), the most common osteoporotic treatment option, stabilize bone mineral through their high binding affinity to hydroxyapatite and decrease bone resorption by suppressing osteoclast activity [13]. BiPs act as non-hydrolyzable analogs of inorganic pyrophosphate, a common calcification inhibitor of ectopic calcification [13, 16, 17]. However, BiPs have demonstrated potential off-target effects on cardiovascular morbidity. BiP usage correlates with higher risk of cardiovascular/cerebrovascular events and arterial fibrillation in osteoporotic patients with a history of previous cardiovascular events [18, 19]. Retrospective clinical trials found paradoxical cardiovascular outcomes in patients taking BiPs. In women with chronic kidney disease, BiPs moderately reduced the hazard ratio of future cardiovascular events in patients with no previous cardiovascular events; yet, in patients with prior cardiovascular events, BiPs increased the hazard ratio of future events [20].

BiPs may affect vascular calcification through both direct alterations to plaque mineral and systemic effects on calcification mediators. Previous studies reported accumulation of BiPs in the arterial wall of both an atherosclerotic rabbit model and human plaques [21, 22]. Short-term BiP treatment elevated serum levels of parathyroid hormone and osteocalcin, key regulators of cardiovascular calcification [23, 24], in a female renal failure rat model [25]. BiPs prevented medial calcinosis in a male renal failure rat model; however, it was parallel to reduction in bone mass density [26]. BiPs altered the mineral microstructure in Apolipoprotein E-deficient (Apoe^-/-^) mice fed an atherogenic diet, indicating a direct interaction between BiPs and mineral within the plaque [1]. Given the long term administration of BiPs for osteoporosis treatment [27], further studies are needed to assess how BiPs affect vascular calcification over the course of plaque development.

Here, we investigated the effect of BiP treatment on atherosclerotic plaque calcification when given at early, intermediate, and late stages of atherosclerosis. We began BiP administration to Apoe^-/-^ mice at three different periods of plaque development based on previous reports [2, 28, 29]: before mineral formation (early), during plaque remodeling and mineral deposition (mid-term), or after the formation of mature, calcified plaques (late stage). Our data indicate that beginning BiP treatments in either early or late stages promote plaque calcification; however, early BiP treatment resulted in narrower mineral size distribution compared to the late-stage treatment. Interestingly, beginning the BiP treatment during active plaque remodeling led to lower levels of plaque calcification compared to the other treatment groups. These data may provide insight into potential cardiovascular-related off-target effects of BiPs.

## 2. Methods

### 2.1. In vivo Study

The animal study was approved by the Institutional Animal Care and Use Committee (IACUC) at Florida International University under protocol 17-022 and conformed to current NIH guidelines. Eight-week-old mice homozygous genotype for the Apoe^tm1Unc^ mutation (B6.129P2-Apoe^tm1Unc^/J, ApoE^-/-^, *n* = 40, 20 per biological sex) were purchased from the Jackson Laboratory (Bar Harbor, Maine, USA). After two weeks of acclimatization and feeding with a regular chow diet, the animals (10 weeks old) were fed with an atherogenic diet (ENVIGO, TD.88137 adjusted calories diet 42% from fat) for 25 weeks. Mice were randomly divided into four groups (5 males and females per treatment): Week 5, Week 10, and Week 15, in which animals received subcutaneous injections of bisphosphonate ibandronate sodium (2 mg/kg/mouse, APEXBIO, B1772) twice per week (on Mondays and Thursdays), following 5, 10, and 15 weeks of the atherogenic diet, respectively. The control group received drug vehicle, phosphate-buffered saline (PBS, 1X, Caisson Labs, PBL01). For the injections, animals were partially anesthetized using isoflurane (1%, Patterson Veterinary, 07-893-1389, in 2 L.min^-1^ oxygen flow). At the study end point (after 25 weeks of diet), the mice received a tail vein injection of a calcium tracer, OsteoSense 680EX (80 nmol/kg/mouse, PerkinElmer, NEV10020EX). After 48 hours, the animals were anesthetized with isoflurane (1%, in 2 L.min^-1^ oxygen flow) followed by retro-orbital bleeding for blood collection. Mice were then immediately euthanized by laceration of the diaphragm before tissue collection. The hearts and aortas were resected and fixed in formalin (10% w/v, Fisher Scientific, SF100) for 2 hours. The blood samples were centrifuged at 2,000×g for 15 minutes to collect the serum.

H2>2.2. Calcification Morphological Quantification

After resection and removal of excess adventitia and adipose tissues, the aortas were imaged using a near-infrared scanner (LI-COR Odyssey) to visualize the atherosclerotic plaque calcification burden. The signals were localized and quantified using edge detection algorithms in a custom MATLAB script, which quantified the total area of the calcium tracer, and normalized to the total scanned aorta area. The resected aortic roots were embedded using Tissue-Plus OCT (Fisher Scientific, 23-730-571) and cryosectioned (sagittal, 18 μm/section). The serial sections were imaged using a confocal microscopy system (Eclipse Ti, Nikon). To assess total, mean, and maximum calcification area in the aortic roots, a custom image analysis script was developed in MATLAB. After filtering the background and smoothing the images, individual microcalcifications were identified as connected pixels in binarized images. The script reported the summation of the pixels, average area of the connected pixels (representing mean of the mineral size), and the maximum connected area (representing size of a single largest mineral) for each image as total, mean, and maximum calcification area, respectively. Connected areas smaller than 5 pixels were excluded from analyzed data.

### 2.3. Serum Alkaline Phosphatase Activity and Total Cholesterol Assessment

To assess the activity of mouse serum tissue non-specific alkaline phosphatase (TNAP), a colorimetric assay kit (BioVision, K412) was used. The samples were diluted 1:20 in assay buffer, and 80 μL of each sample were mixed with 50 μL of 5 mM pNPP solution and incubated for 60 min at 25°C. The colorimetric change resulting from the reaction was detected using a multi-mode reader (BioTek, Synergy HTX) to measure absorbance at 405 nm.

To quantify the serum total cholesterol, a Wako Free Cholesterol E kit (FUJIFILM Medical Systems USA, 99902601) was used. Briefly, 10 μL of each serum sample were resuspended in 1 mL of assay color reagent and incubated at 37°C for 5 min. The colorimetric change resulting from the reaction was detected using a plate reader to measure absorbance at 600 nm.

### 2.4. X-ray Computed Tomography (X-ray CT)

Femurs were dissected from mice and imaged directly in a Nikon XT H 225 scanner (macro-CT, Nikon Metrology, Tring, UK). The raw transmission images were reconstructed using commercial image reconstruction software package (CT Pro 3D, Nikon Metrology, Tring, UK), which employs a filtered back-projection algorithm. The scan was performed using 80 □kV beam energy, 70 □μA beam current, and a power of 5.6 W. A PerkinElmer 1620 flat panel detector was used, with 200 □μm pixel size. The resulting effective pixel size was 5 □μm. The exposure time per projection was 0.5 s, and a total of 1601 projections were acquired, resulting in a scanning time of approximately 13 minutes per sample. Bone structural parameters, including thickness and volume fraction (the ratio of bone volume (BV) to total volume (TV)), for both cortical and trabecular regions were assessed using a plug-in module, BoneJ [30], in ImageJ (NIH, USA) [31].

### 2.5. Statistics

Data are presented as the mean of independent replications, and error bars represent the standard error of the mean. The reported *n* values represent independent biological replicates. Statistical significance between groups was calculated using two-way ANOVA with mixed-effects model in GraphPad Prism 8. A p-value less than 0.05 was considered statistically significant. In the case of comparisons between two groups, the statistical significance was calculated using Welch’s t-test with p-values less than 0.05 significant.

## 3. Results

### 3.1. BiP treatment increases atherosclerotic plaque calcification in the Apoe^-/-^ mice fed an atherogenic diet

Visualization and quantification of the calcium tracer OsteoSense from resected aorta revealed significantly elevated plaque calcification in all BiP-treated animals compared to the control group (**Fig. 1, A**), regardless of BiP regimen beginning time and sex. In male mice, early (week 5) and late (week 15) BiP treatments showed the highest plaque calcification compared to control and mid-term BiP administration (week 10), as shown in **Fig. 1, B**. However, female mice exhibited no significant difference among the BiP treated groups in terms of plaque calcification burden along the aorta (**Fig. 1, C**). Interestingly, female mice showed significantly higher levels of calcification compared to males when the BiP treatment began at 10 weeks of the atherogenic diet (Fig. 1, F). The level of calcification was similar between males and females in control, week 5, and week 15 groups (**Fig. 1, D** to **G**). These data demonstrate that BiP may interact with the ectopic atherosclerotic plaque calcification and increase the rate of mineral formation in a sex-dependent manner.

**Figure 1.**
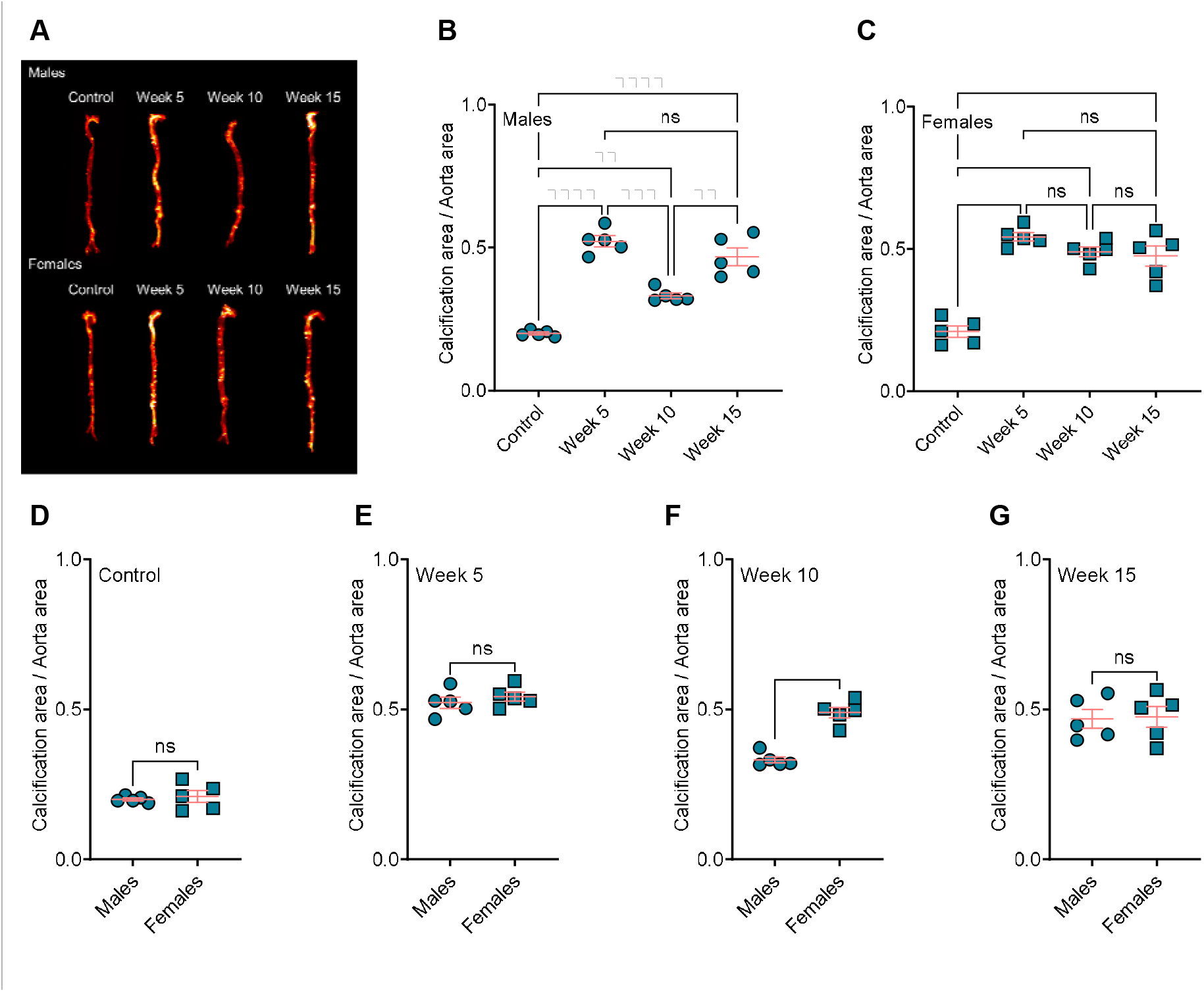
BiP treatment increases the atherosclerotic plaque calcification in Apoe^-/-^ mice fed an atherogenic diet. (A) visualization of the calcium burden using a near-infrared calcium tracer, OsteoSense; (B and C) quantification of OsteoSense signal and correlation to total calcification in male and female mice, respectively; (D to G) comparison of total calcification between male and female mice in each group. **P ≤ 0.01, ***P ≤ 0.001, and ****P ≤ 0.0001, two-way ANOVA with Tukey’s post-hoc test for comparison between multiple groups, and Welch’s t-test for comparison between two groups.

### 3.2. BiP treatment alters the size of mineral in the atherosclerotic plaque

Analysis of the mineral morphologies in the aortic roots of the mice showed BiP treatment significantly increased the total calcification area in both male and female groups, compared to the control mice (**Fig. 2, A, B, E**, and **F**). Similar to the plaque calcification along the aorta, both week 5 and week 15 male groups had a significantly higher level of total calcification area, compared to the week 10 group (**Fig. 2, B**). However, no significant changes were observed between the female BiP-treated groups. The average calcification size was significantly bigger in early BiP-treated group (week 5) compared to the control, week 10, and week 15 groups. The mean calcification area showed no significant differences between the male control, week 10, and week 15 groups (**Fig. 2, C**). In female mice, the mean calcification area increased in both week 5 and week 15, compared to the control and week 10 groups. No significant changes were observed between the control and week 10 groups (**Fig. 2, G**). Maximum calcification area, the largest mineral detected in a plaque, increased in all BiP-treated groups compared to the control, regardless of sex. The largest calcified area in the male groups was detected in both week 5 and week 15. In female groups, early BiP treatment increased the maximum mineral size in the plaque compared to other time points, with no significant differences between week 10 and 15 (**Fig. 2, H**).

**Figure 2.**
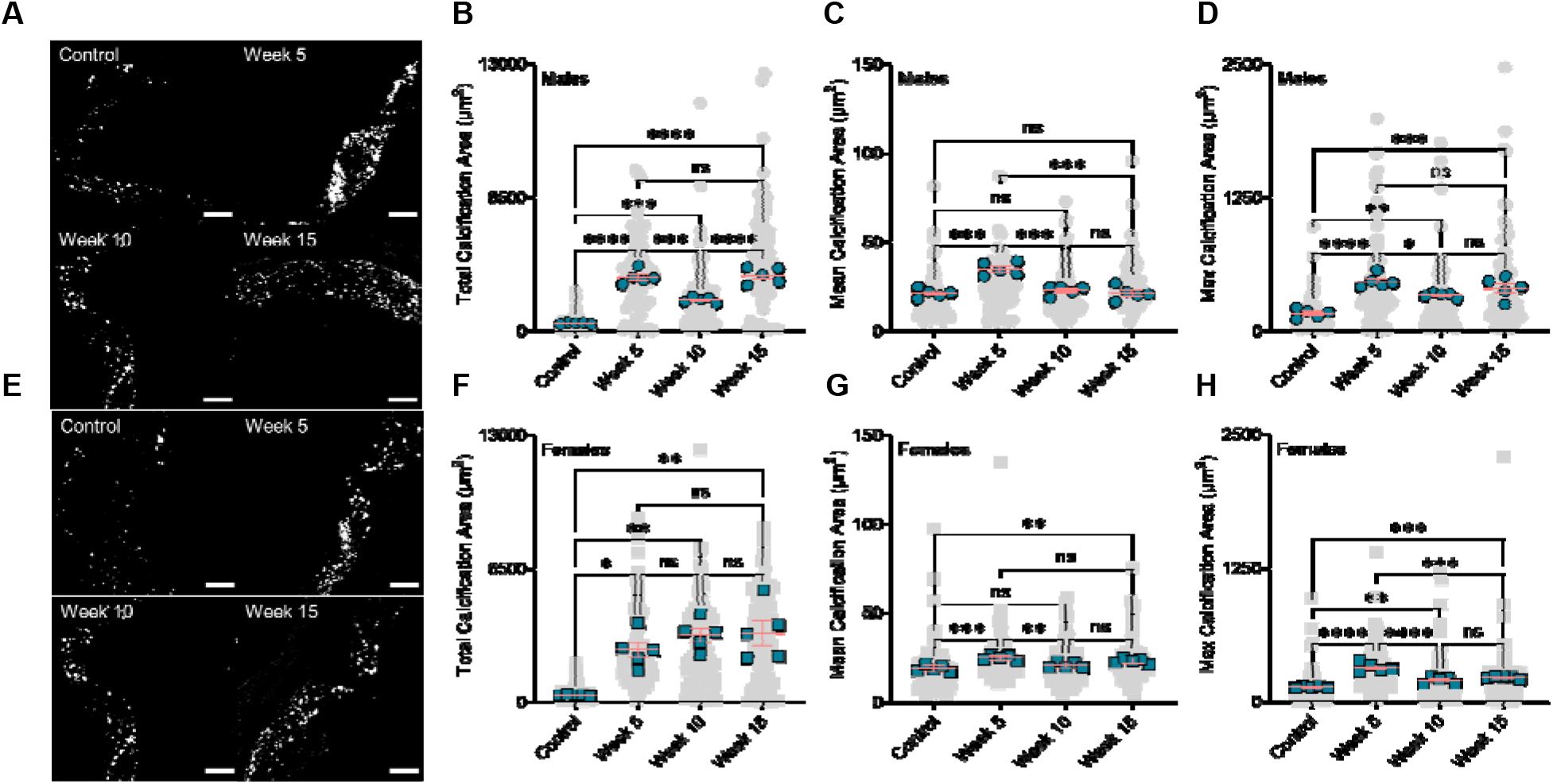
BiP treatment alters the micromorphology of the minerals in the atherosclerotic plaque. (A) visualization of the minerals in the atherosclerotic plaques of male mice (20X, scale bar 100 μm); (B) total plaque calcification in male mice; (C) mean calcification area (mean of mineral size) in male animals; (D) maximum calcification area in male mice; (E) visualization of the minerals in the atherosclerotic plaques of female mice (20X, scale bar 100 μm); (F) total plaque calcification in female mice; (G) mean calcification area (mean of mineral size) in female animals; (H) maximum calcification area in female mice. *P < 0.05, **P ≤ 0.01, ***P ≤ 0.001, and ****P ≤ 0.0001, two-way ANOVA with Tukey’s post-hoc test.

Comparing male and female mice, we observed no significant differences in total calcification area in the aortic roots of control, week 5, and week 15 treatment groups. However, BiP treatment beginning at 10 weeks of diet significantly increased the total calcification area in the female mice compared to the male mice (**Fig. 3**, panel **A**). The mean calcification area (mineral average size) remained unchanged between the males and females in the control, week 10, and week 15; however, in week 5, the male mice showed a significantly elevated mean calcification area compared to the females (**Fig. 3**, panel **B**). Regardless of the time at which BiP treatment began, the maximum calcification area in the male groups was higher than the females; however, no significant differences were detected in the control group between males and females (**Fig. 3**, panel **C**). These data suggest that BiP treatment may interact with and alter the morphology of mineral in the plaque.

**Figure 3.**
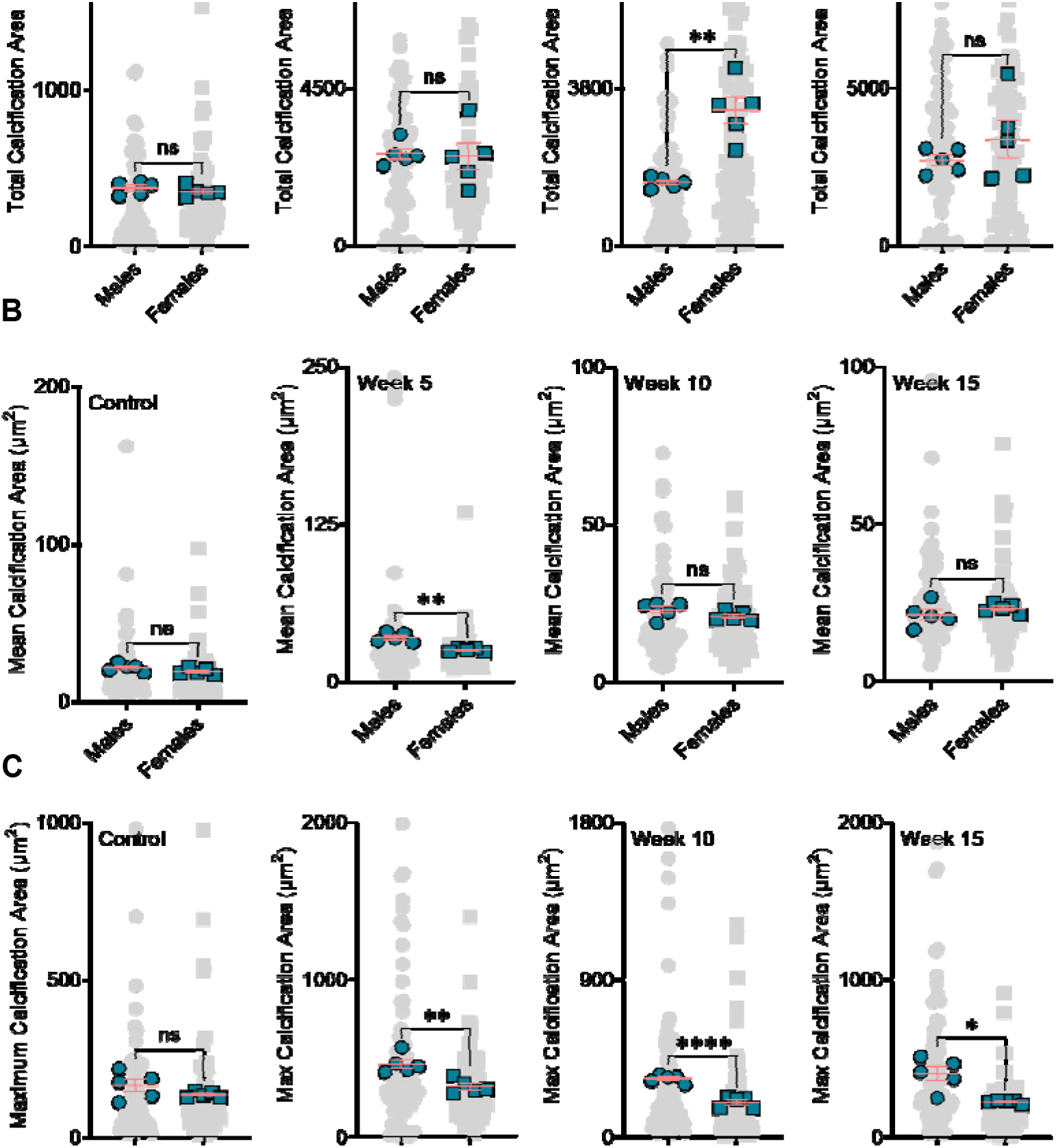
BiP treatment may affect the mineral morphology differently in male and female mouse model. Panel (A) comparison of total plaque calcification between male and female mice; Panel (B) comparison of mean calcification area between male and female mice; Panel (C) comparison of maximum calcification area between male and female mice. *P < 0.05, **P ≤ 0.01, and ****P ≤ 0.0001, Welch’s t-test.

### 3.3. BiP treatment does not affect serum alkaline phosphatase activity and total cholesterol level

We measured the level of alkaline phosphatase, an enzyme required for calcification, in the serum collected at the study endpoint. We observed no significant differences between the BiP-treated and the control groups across either sex (**Fig. 4, A** and **B**). Furthermore, the total serum cholesterol remained unchanged across BiP-treated groups compared to the controls for both male and female mice (**Fig. 4, C** and **D**).

**Figure 4.**
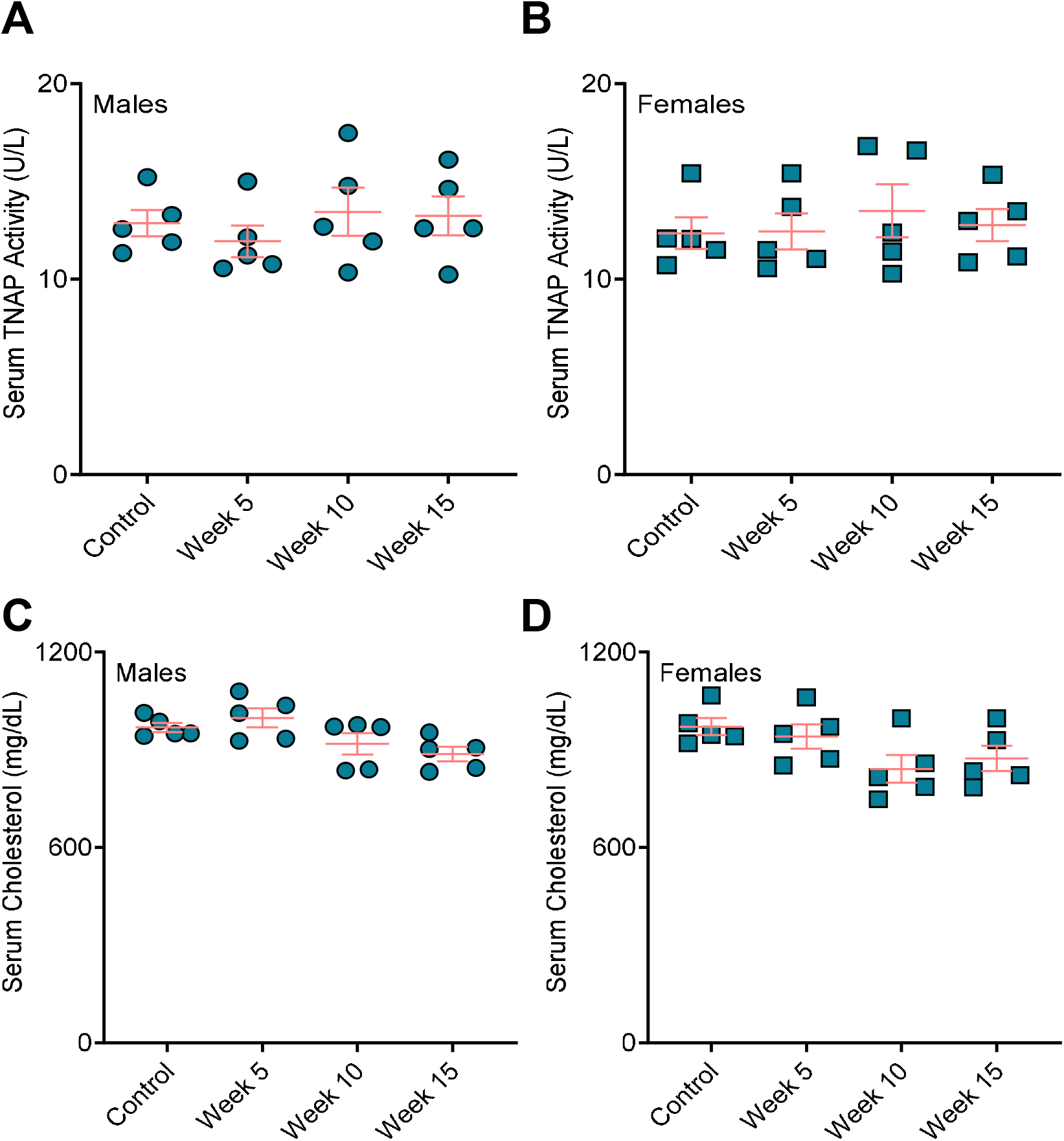
BiP treatment does not affect the serum TNAP activity and total cholesterol in mouse model of atherosclerotic plaque calcification. Serum TNAP activity in (A) male mice and (B) female mice; serum total cholesterol in (C) male mice and (D) female mice. No statistically significance observed across the groups, two-way ANOVA with Tukey’s post-hoc test.

### 3.4. Bone remodeling positively correlates with the BiP treatment duration

Analyses of the resected bones showed dramatic increased bone volume in BiP-treated mice compared to the control group. The longer treatment with BiP (i.e., beginning the BiP treatment at the early time point) results in a higher bone volume in both cortical and trabecular areas (metaphyseal and epiphyseal regions), as shown in **Fig. 5**, panel **A** and **B**. The cortical bone thickness followed a similar trend and exhibited a positive correlation with the duration of BiP treatment. The thickness of the trabecular bone, both metaphyseal and epiphyseal regions, was significantly increased in the longest BiP-treated group (beginning of BiP regimen after 5 weeks of diet); this parameter remained unchanged for week 10 and week 15 groups compared to the control mice. **Table 1** summarizes the bone microstructure parameters analyzed for each group.

**Figure 5.**
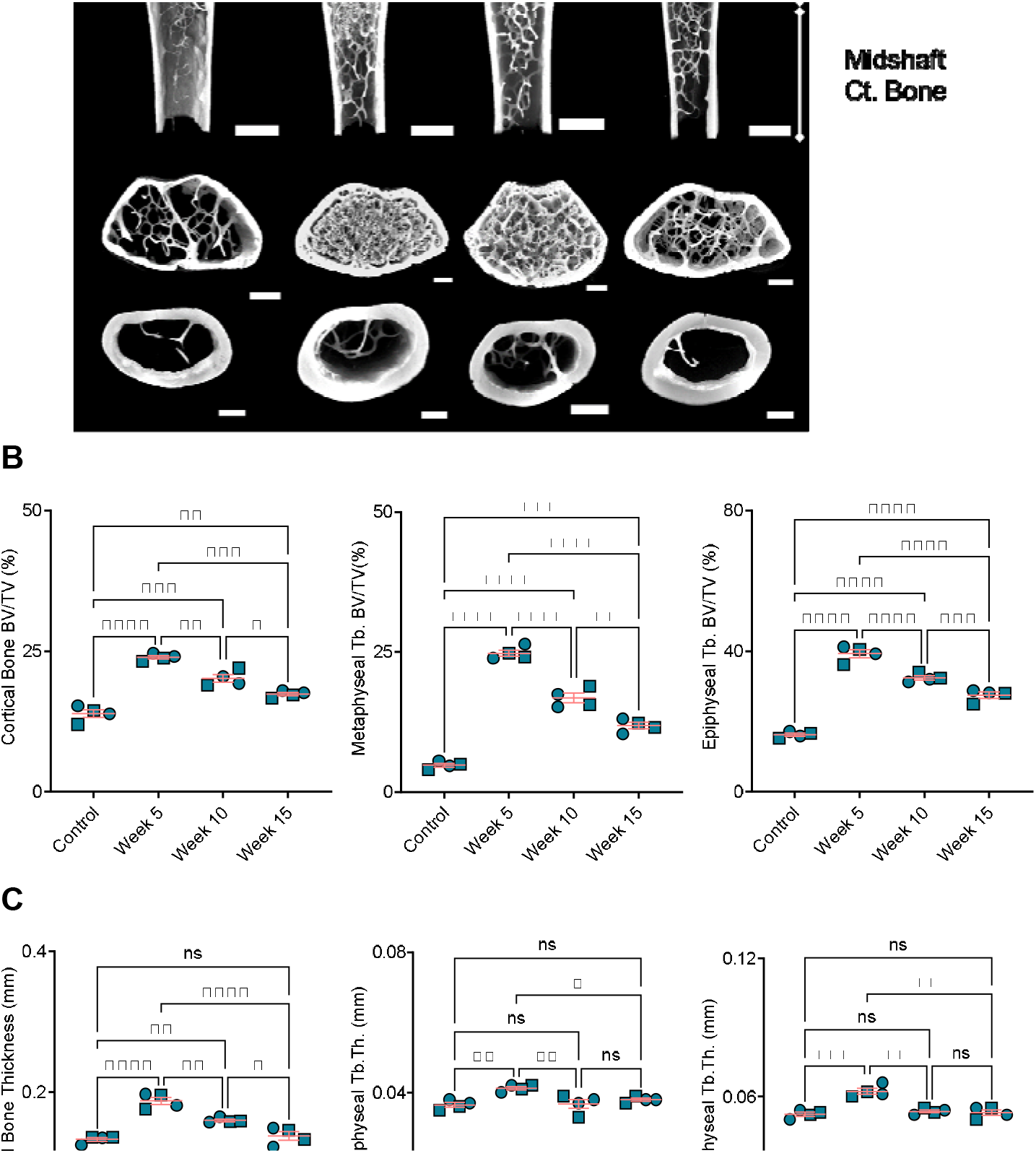
Bone remodeling positively correlates with duration of BiP treatment. (A) reconstruction of femoral bone microstructure for the different treated and untreated groups; Panel (B) bone volume fraction (bone volume/total volume) for cortical and trabecular regions; Panel (C) bone thickness for cortical and trabecular regions. *P < 0.05, **P ≤ 0.01, ***P ≤ 0.001, and ****P ≤ 0.0001, two-way ANOVA with Tukey’s post-hoc test. Male mice (•) and female mice (◼).

**Table 1.** Detailed bone structural parameters.

| Parameter | Control<br>*n* = 4 | Week 5<br>*n* = 4 | Week 10<br>*n* = 4 | Week 15<br>*n* = 4 |
| --- | --- | --- | --- | --- |
| **Distal femur** |  |  |  |  |
| *Epiphyseal Trabecular Bone* |  |  |  |  |
| BV/TV (%) | 16.22 ± 0.37 | 39.3 ± 0.96 | 32.4 ± 0.5 | 27.48 ± 0.76 |
| Tb.N (mm<sup>-1</sup>) | 7.63 ± 1.11 | 12.86 ± 0.11 | 12.82 ± 0.32 | 12.05 ± 0.34 |
| Tb.Th (mm) | 0.052 ± 0.001 | 0.062 ± 0.001 | 0.053 ± 0.001 | 0.053 ± 0.001 |
| Tb.Sp (mm) | 0.59 ± 0.06 | 0.41 ± 0.02 | 0.42 ± 0.02 | 0.43 ± 0.02 |
| BS/BV<br>(mm<sup>2</sup>/mm<sup>3</sup>) | 0.065 ± 0.002 | 0.059 ± 0.003 | 0.057 ± 0.003 | 0.062 ± 0.003 |
| EF | 0.017 ± 0.003 | 0.033 ± 0.005 | 0.034 ± 0.009 | 0.016 ± 0.003 |
| EF<sub>max</sub> | 0.885 ± 0.004 | 0.902 ± 0.005 | 0.887 ± 0.004 | 0.88 ± 0.009 |
| EF<sub>min</sub> | -0.82 ± 0.003 | -0.825 ± 0.008 | -0.812 ± 0.007 | -0.81 ± 0.009 |
| DA | 1.43 ± 0.05 | 1.39 ± 0.04 | 1.42 ± 0.03 | 1.4 ± 0.07 |
| *Metaphyseal Trabecular Bone* |  |  |  |  |
| BV/TV (%) | 4.82 ± 0.29 | 24.8 ± 0.5 | 16.8 ± 0.75 | 11.86 ± 0.5 |
| Tb.N (mm<sup>-1</sup>) | 4.1 ± 0.3 | 13.1 ± 0.09 | 11.02 ± 0.07 | 7.62 ± 0.49 |
| Tb.Th (mm) | 0.036 ± 0.001 | 0.041 ± 0.001 | 0.037 ± 0.001 | 0.038 ± 0.001 |
| Tb.Sp (mm) | 0.56 ± 0.094 | 0.29 ± 0.04 | 0.27 ± 0.015 | 0.44 ± 0.022 |
| BS/BV<br>(mm<sup>2</sup>/mm<sup>3</sup>) | 0.092 ± 0.003 | 0.085 ± 0.001 | 0.092 ± 0.003 | 0.092 ± 0.01 |
| EF | 0.12 ± 0.02 | 0.07 ± 0.005 | 0.086 ± 0.017 | 0.07 ± 0.03 |
| EF<sub>max</sub> | 0.835 ± 0.019 | 0.87 ± 0.002 | 0.86 ± 0.012 | 0.855 ± 0.03 |
| EF<sub>min</sub> | -0.67 ± 0.036 | -0.765 ± 0.013 | -0.71 ± 0.022 | -0.73 ± 0.05 |
| DA | 1.77 ± 0.14 | 1.6 ± 0.15 | 1.522 ± 0.09 | 1.78 ± 0.09 |
| **Femoral midshaft** |  |  |  |  |
| *Cortical Bone* |  |  |  |  |
| Mean Ct.Th (mm) | 0.133 ± 0.002 | 0.187 ± 0.004 | 0.159 ± 0.001 | 0.137 ± 0.005 |
| BV/TV (%) | 13.9 ± 0.48 | 23.9 ± 0.25 | 20.2 ± 0.6 | 17.3 ± 0.24 |
| Ps.Pm (mm) | 5.65 ± 0.11 | 5.37 ± 0.2 | 5.35 ± 0.22 | 5.33 ± 0.18 |
| Ec.Pm (mm) | 4.75 ± 0.11 | 4.2 ± 0.18 | 4.15 ± 0.2 | 4.17 ± 0.21 |
| J (mm<sup>4</sup>) | 0.49 ± 0.03 | 0.455 ± 0.04 | 0.417 ± 0.007 | 0.5 ± 0.1 |

Data were combination of males and females, and collected using X-ray Computed Tomography of distal femur and femoral midshaft. Values represent the mean ± SEM.
BV/TV = Bone Volume/Total Volume; Tb.N = Trabecular Number; Tb.Th = Trabecular Thickness; Tb.Sp = Trabecular Separation; BS/BV = Specific Bone Surface: Bone Surface Area/Bone Volume; EF = Ellipsoidal Factor; DA = Degree of Anisotropy; Ct.Th = Cortical Thickness; Ps.Pm = Periosteal Perimeter; Ec.Pm = Endocortical Perimeter; J = Polar Moment of Inertia.

## 4. Discussion

Atherosclerotic plaque calcification in Apoe^-/-^ mice fed a chow diet begins around 45-60 weeks of age [32]; however, feeding the mice an atherogenic diet (21% fat and 15% cholesterol) accelerates plaque calcification dramatically following 10-12 weeks of the diet [29, 33]. Apoe^-/-^ mice fed an atherogenic diet develop early plaque, including lipoproteins accumulation, immune system activation, and formation of cholesterol-rich foam cells between 4 to 14 weeks of feeding. Lipid cores develop following 14 to 16 weeks of the diet, and fibrous cap formation and mature plaque development occur over 18 to 20 weeks of the diet [34]. Here, we studied the role of BiP treatment on atherosclerotic plaque calcification in Apoe^-/-^ mice. The animals received BiP treatment twice per week at three different time points: after 5 weeks of the diet, when the plaque formation has started but prior to calcification formation; after 10 weeks of diet, at the onset of plaque calcification; and after 15 weeks of diet, when the plaque is developed and contains calcified regions.

We showed that the total calcification burden was increased by both the early (starting after 5 weeks of diet and continued for 20 weeks) and late (starting after 15 weeks of diet and continued for 10 weeks) BiP treatment regimens in male mice compared to the control group that did not receive BiP treatment. The early BiP treatment led to a higher mean calcification area (bigger average mineral size) compared to the late treatment, which may stabilize the plaque by reducing the rupture risk. Given the fact that maximum calcification area was comparable between early and late BiP treatment groups, a smaller mineral size average in the week 15 group means the presence of a wider range of mineral size distribution, which may influence the plaque stability. Beginning BiP treatment at week 10 resulted in the smallest average and maximum mineral size compared to other BiP treated male mice, however not significantly different from week 15. The female mice showed the same trends regarding average and maximum calcification size, with the smallest average mineral observed in the mice that received BiP treatment beginning at week 10. However, the differences between treatment groups were less pronounced in the female mice. These data suggest that treating mice before calcification begins (week 5) results in the formation of larger calcifications, whereas starting BiP treatment during early (week 10) or late (week 15) stages of pro-calcific plaque remodeling leads to smaller average mineral size.

Mouse atherosclerotic plaques do not rupture, which limited our ability to perform plaque stability analyses in this study. However, our data suggest that timing of BiP intervention in relation to ongoing atherosclerotic remodeling could influence mineral morphology and thus plaque stability. Previous studies using electron microscopy revealed that timing of BiP regimen affects the morphology and topography of plaque microcalcifications (less than 5 μm^2^) [1]; starting BiP treatment in early plaque remodeling led to bigger individual mineral aggregates with higher surface roughness, while starting at later time points reduced both mineral aggregates size and surface roughness [1]. Furthermore, mineral aggregates associated with later BiP treatment beginning showed qualitatively disorganized and loose morphologies [1]. Finite element-based predictions of these previous data suggested that the microcalcifications associated with all BiP treatment groups would reduce plaque stress compared to non-treated controls. The present data provide a more global analysis of calcification size and burden following BiP treatment and support the previous observation that BiP alters calcification morphology.

We showed that starting BiP treatment at early and late stages of calcific remodeling affected the total calcification area similarly in male and female mice. However, for the BiP regimen given during plaque remodeling and mineral deposition onset, total calcification area was higher in female mice compared to males. The mean calcification area (average of the mineral size) was similar across males and females in this group; yet, male mice had a larger maximum calcification area compared to the females, which suggest a wider mineral size distribution in males. These results may indicate a critical timing in respect of BiP usage. Recent clinical studies indicate a negative correlation between the cumulative BiP dosage (an indication of portion of day covered) to hazard ratio of hospitalization due to atherosclerotic cardiovascular events [35]. Female patients indicated higher hazard ratio (0.958) compared to associated male patients (0.897) [35], an effect that may explain the sex-dependent observation in our week 10 group. Furthermore, these clinical outcomes support our data that the longer BiP usage duration may lead to larger minerals with a narrow size distribution, which *in silico* studies suggest associate with stable plaque and lower rupture risk [8, 36].

In humans, 95% of the serum TNAP originate from bone and liver [37], and unbalanced bone turnover, aging, and chronic kidney disease correlate with abnormal elevated serum TNAP [38, 39]. Serum TNAP increases in osteoporotic postmenopausal women compared to postmenopausal women without osteoporosis [38]. BiPs lowered serum TNAP in osteoporotic postmenopausal women and men with heterotopic ossification [38, 40, 41]. Our data showed unchanged TNAP in serum across the treated and untreated groups. Thus, the observed changes in plaque calcification in all treated groups appears to be independent of systemic TNAP changes.

Clinical trials reported serum LDL reduction in osteoporotic females treated with BiPs; however, the level of high-density lipoprotein and lactate dehydrogenase remained unchanged [42-44]. In the present study, serum total cholesterol remained unchanged in all animals, regardless of the BiP treatment. Our data indicate that the observed calcification differences by BiP treatment are not due to effects on lipid metabolism in Apoe^-/-^ mice.

The outcomes from bone microstructure analyses demonstrate the substantial effect of BiP treatment on bone remodeling. These results reveal a positive correlation between BiP regimen treatment duration and the bone volume and thickness in cortical and trabecular areas. We showed a positive correlation between bone remodeling and increased plaque calcification for early BiP treatment in Apoe^-/-^ mice. Long term BiP treatment led to increased mineralization in both bone and atherosclerotic plaque. However, for later time points, beginning BiP treatment after 10 and 15 weeks of diet, the data demonstrate a negative correlation between bone remodeling and plaque calcification. Both cortical and trabecular bone volumes are significantly higher in the week 10 group, compared to week 15, while plaque calcification for the latter is significantly higher compared to former. Previous studies reported that BiP may imitate the effects of pyrophosphate, a mineralization inhibitor in the cardiovascular system; however, our data did not support the inhibitory role of BiP to reduce cardiovascular calcification compared to non-treated controls. Importantly, the choice of animal models or patients in clinical trials may affect the BiP outcomes; atherosclerotic plaque calcification represents a complex process in which several cell types, including VSMCs and macrophages [45, 46], are involved, thus, the BiP treatment may affect multiple aspects of plaque progression and calcification.

## 5. Conclusion

Atherosclerotic plaque calcification represents a significant predictor of lesion vulnerability. Patients with bone disorders are prone to develop ectopic cardiovascular calcification. Clinical trials correlated BiPs, a common osteoporotic pharmaceutical family, with contradicting cardiovascular outcomes. Here, we demonstrated the importance of treatment timing in BiP-induced mineral disruption or promotion. We indicated that BiP can alter key morphological features of the microcalcifications within the atherosclerotic plaque of Apoe^-/-^ mice, which may determine the risk of plaque rupture. Early beginning of BiP regimen, i.e., before calcification initiates, increased the total calcification in both males and females; however, the treatment led to narrower mineral size distribution. Later regimen timings, either after mineralization starts or when the plaque is developed, result in wider mineral size distribution, which correlates with plaque destabilization and a higher risk of rupture. Interestingly, early BiP regimen elevated mineralization in both bone and atherosclerotic plaque. However, BiP regimen correlates negatively with bone mineralization and plaque calcification if given after the onset of plaque mineralization.

## Author Contributions

The author contributions are as following: Conceptualization, A.B.N. and J.D.H.; Methodology, S.B. and J.D.H.; Software, A.B.N., M.G.R. and J.D.H.; Validation, A.B.N., M.G.R., F.I., P.R.S., S.B. and J.D.H.; Formal Analysis, A.B.N., F.I., P.R.S. and J.D.H.; Investigation, A.B.N., H.H.N., F.I. and P.R.S.; Resources, H.H.N., F.I. and J.D.H.; Data Curation, A.B.N. and M.G.R.; Writing – Original Draft Preparation, A.B.N.; Writing – Review & Editing, H.H.N., M.G.R., F.I., P.R.S., S.B. and J.D.H.; Visualization, A.B.N.; Supervision, J.D.H.; Project Administration, J.D.H.; Funding Acquisition, J.D.H.

